# Periaqueductal efferents to dopamine and GABA neurons of the VTA

**DOI:** 10.1101/117416

**Authors:** Niels R. Ntamati, Meaghan Creed, Christian Lüscher

## Abstract

Neurons in the periaqueductal gray (PAG) modulate threat responses and nociception. Activity in the ventral tegmental area (VTA) on the other hand can cause reinforcement and aversion. While in many situations these behaviors are related, the anatomical substrate of a crosstalk between the PAG and VTA remains poorly understood. Here we describe the anatomical and electrophysiological organization of the VTA-projecting PAG neurons. Using rabies-based, cell type-specific retrograde tracing, we observed that PAG to VTA projection neurons are evenly distributed along the rostro-caudal axis of the PAG, but concentrated in its posterior and ventrolateral segments. Optogenetic projection targeting demonstrated that the PAG-to-VTA pathway is predominantly excitatory and targets similar proportions of *I*_h_-expressing VTA DA and GABA neurons. Taken together, these results set the framework for functional analysis of the interplay between PAG and VTA in the regulation of reward and aversion.

## Introduction

The periaqueductal gray (PAG) is a heterogeneous midbrain structure that is critical for the endogenous modulation of nociception and for the expression of defensive behaviors [1]. These functions have been shown to be mediated by neurons anatomically segregated in the longitudinal columns corresponding to the dorsolateral (dl), lateral (l) and ventrolateral (vl) subdivisions of the PAG [2]. The PAG is also the major site of action for opioid analgesia [3,4].

Anatomical tracing studies have described ascending and descending projections from the PAG to a variety of brain structures [5]. Among these projection targets is the ventral tegmental area (VTA), a major component of the brain reward system [6]. While it is primarily known for its role in reward prediction, and positive reinforcement [7,8], the VTA has also been implicated in the modulation of nociception and in the expression of fear and aversive responses [9–12]. The engagement of VTA dopamine- (DA) and *gamma*-aminobutyric acid (GABA)-releasing neurons with PAG afferents through both symmetric and asymmetric synaptic contacts has been demonstrated with rabies-assisted retrograde tracing and ultrastructural immunoelectron microscopic analyses [13,14].

It remains elusive, however, whether the input neurons providing these VTA afferents spatially segregate within specific PAG columns, potentially associating the PAG-to-VTA pathway with specific anti-nociceptive or defensive functions. Moreover, evidence for an electrophysiologically functional connection, and knowledge of its excitatory or inhibitory effect onto VTA DA and GABA neurons is still lacking. To this end, the present study will describe the rostro-caudal distribution of VTA-projecting neurons across the PAG columns, and will test whether these neurons exhibit a preferential excitatory or inhibitory effect on DA and GABA neurons of the VTA.

## Materials and methods

### Animals

Experiments were performed on DAT-Cre [15] and GAD65-Cre mice [16] of both sexes. All animal procedures were performed in accordance with the authors' university animal care committee's regulations.

### Injection procedures

All stereotaxic intracranial injections were performed under isoflurane anesthesia (2-5%, Attane) using glass capillary pipettes connected to a microinjection pump (Narishige) at a rate of ~100 nl min^−1^. The coordinates used were (from bregma, in mm): AP −3.4, ML ±0.5, DV −4.3 for VTA injections; AP −4.0, ML ±0.3, DV −2.6 for PAG injections. For retrograde tracing experiments, 300 nl of a 1:1 mixture of AAV8-CAG-DIO-RG and AAV5-EF1a-DIO-TVA-mCherry was injected unilaterally in the VTA, followed 2 weeks later by the injection of 800 nl of RV ΔG-EnvA-EGFP at the same coordinates. For patch clamp experiments, animals were bilaterally injected with AAV5-EF1a-DIO-mCherry in the VTA and with AAV2-hsyn-ChR2-EYFP in the PAG (all viruses from the UNC Vector Core Facility).

### Electrophysiology on acute slices

Coronal VTA slices (180 μm) were prepared using a vibratome in ice-cold cutting solution containing (in mM): NaCl 87, NaHCO_3_ 25, KCl 2.5, MgCl_2_ 7, NaH_2_PO_4_ 1.25, CaCl_2_ 0.5, glucose 25, and sucrose 75. Slices were incubated in the same solution for 20 min at 31°C before being transferred to regular room-temperature artificial cerebrospinal fluid (aCSF), containing (in mM): NaCl 119, NaHCO_3_ 26.2, KCl 2.5, MgCl_2_ 1.3, NaH_2_PO_4_ 1, CaCl_2_ 2.5, and glucose 11. After at least one hour for recovery, slices were transferred to the recording chamber, superfused with aCSF at 2 ml/min. All solutions were constantly bubbled with 95% O_2_ and 5% CO_2_. Postsynaptic currents were evoked by stimulating ChR2 with brief (4 ms) blue light pulses using a 470nm LED mounted on the microscope and powered by an LED driver under computer control. Kynurenic acid (kyn, 4 mM, Sigma) and picrotoxin (PTX, 200 μM, Sigma) were added to the aCSF in order to block glutamate- and GABA-mediated currents, respectively. All representative traces were made from averaging at least 20 consecutive sweeps. Neurons were visually identified using an IR CCD camera mounted on an Olympus BX45 microscope. Borosilicate glass pipettes at a resistance range of 2-4 MΩ were used for recording. The internal solution used contained (in mM): K-gluconate 30, KCl 100, MgCl_2_ 4, creatine phosphate 10, Na_2_ATP 3.4, Na_3_GTP 0.1, EGTA 1.1, and HEPES 5. The calculated reversal potential with this internal solution for Cl^−^ was −5mV and cells were voltage-clamped at −70 mV. Currents were amplified, filtered at 2 kHz, digitized at 10 kHz, and saved on a hard disk. The liquid junction potential was small (−4 mV) and traces were therefore not corrected. Access resistance was monitored by a hyperpolarizing step of −4 mV at the onset of every sweep, and the experiment was discarded if the access resistance changed by more than 20%.

### Histological procedures and imaging

Mice were deeply anesthetized with pentobarbital (300 mg/kg i.p.) and transcardially perfused with 0.1M phosphate buffered saline (PBS) followed by 4% paraformaldehyde (PFA, Sigma). Brains were removed, post-fixed in 4% PFA for 24 h at 4°C, and cut in 50 μm sections on a vibratome. For tyrosine hydroxylase (TH) immunohistochemistry, brain sections were rinsed in PBS (0.1 M) and incubated for 1 h at room temperature in a blocking solution containing 5% bovine serum albumin (Sigma) and 0.3% Triton X-100 (Axon Lab AG) in PBS. Sections were then incubated overnight at +4° C with a rabbit anti-TH antibody (1:500; Millipore) in blocking solution, then rinsed in PBS, incubated for 2 h at room temperature with an alexa-conjugated goat anti-rabbit antibody (1:500; Invitrogen) in blocking solution. Sections were then rinsed again in PBS and mounted on glass slides with Fluoroshield Mounting Medium (Abcam). Confocal images were captured with a Nikon A1r Spectral scanning confocal microscope, and then processed with ImageJ.

### Data analysis and statistics

Equal regions of interest (ROIs) were selected for rabies-infected starter cell counts and then averaged among all mice. Cell counts of retrogradely labeled input neurons in the PAG were performed using Paxinos and Franklin's mouse brain atlas as reference to divide the PAG into dorsolateral, lateral and ventrolateral segments, and into rostral (AP coordinates between −2.8 and −3.4), central (between −3.4 and −4.16) and caudal regions (between −4.16 to −5.0) [17]. All data are expressed as mean ± standard error of the mean, and statistical analysis was performed with Igor, Microsoft Excel, and GraphPad Prism.

## Results

### PAG neurons project to VTA DA and GABA neurons

To reveal the distribution of PAG to VTA projecting neurons, we employed a cell-type specific, virally assisted tracing approach using a rabies virus pseudotyped with the avian sarcoma and leucosis virus envelope glycoprotein EnvA and lacking the rabies envelope glycoprotein gene *rabG* (RVΔG-EGFP, [18]). First, we virally transfected the VTA of DAT-Cre or GAD65-Cre mice with two Cre-dependent constructs encoding for the EnvA receptor TVA and RabG (DIO-TVA-mCherry/RG), and followed up with the RVΔG-EGFP transfection two weeks later (Fig. 1A). Seven days later, infection of VTA DA and GABA neurons and their retrogradely labeled afferents were observed (Fig. 1B). VTA neurons co-expressing RVΔG-EGFP and the DIO-TVA-mCherry/RG constructs were identified as starter cells, from which the inputs were monosynaptically tagged by the expression of RVΔG-EGFP alone (Fig. 1C). We then visualized the cell bodies of these VTA projection neurons in the PAG. Given its extensive rostro-caudal extension, we partitioned the PAG into three segments that represent the columnar divisions as described in Paxinos and Franklin's mouse brain atlas [17]. The rostral segment includes the PAG from the anteroposterior (AP) coordinates −2.8 and −3.4 (in mm, from bregma). The central segment, subdivided in dorsolateral PAG (dlPAG) and lateral PAG (lPAG), lies between AP −3.4 and −4.16. The caudal PAG, subdivided in dlPAG, lPAG and ventrolateral PAG (vlPAG), extends from AP −4.16 to −5.0 (Fig. 1D). We observed that the PAG projection neurons were sparse in the rostral PAG, and increasingly more dense in the central and caudal PAG segments. We found that VTA DA- and GABA-projecting neurons are similarly distributed along the PAG, with the exception for the caudal PAG, where more inputs are provided to VTA GABA neurons (AP coordinate −4.6, DA vs GABA: 17.3 ± 4.4 vs 53.8 ± 11.3), despite a similar number of starter cells compared to DA neurons (DA vs GABA: 96.5 ± 18.3 vs 92.7 ± 5.6) (Fig. 1E). We then analyzed the intracolumnar distribution of VTA projecting neurons (Fig. 1F). We found that both VTA DA and GABA inputs are similarly scattered across the PAG columns, with the caudal ventrolateral and central lateral ones providing the highest proportion of total PAG inputs (DA: caudal vlPAG, 38.8 ± 6.9, central lPAG, 31.3 ± 2.4; GABA: caudal vlPAG, 33.3 ± 3.4, central lPAG, 32.7 ±3.9). While some inputs were observed to decussate from the contralateral side, the majority DA and GABA inputs was similarly labeled in the ipsilateral hemisphere, indicating a moderate level of input lateralization (DA vs GABA: 0.77 ± 0.03 vs 0.69 ± 0.02). Taken together, these results indicate that both VTA DA- and GABA-projecting neurons from the PAG concentrate ipsilaterally and preferentially in its lateral and ventrolateral subdivisions, in stark contrast with the dorsolateral columns which provide less than 5% of the total PAG inputs (Fig. 1G).

**Figure 1.**
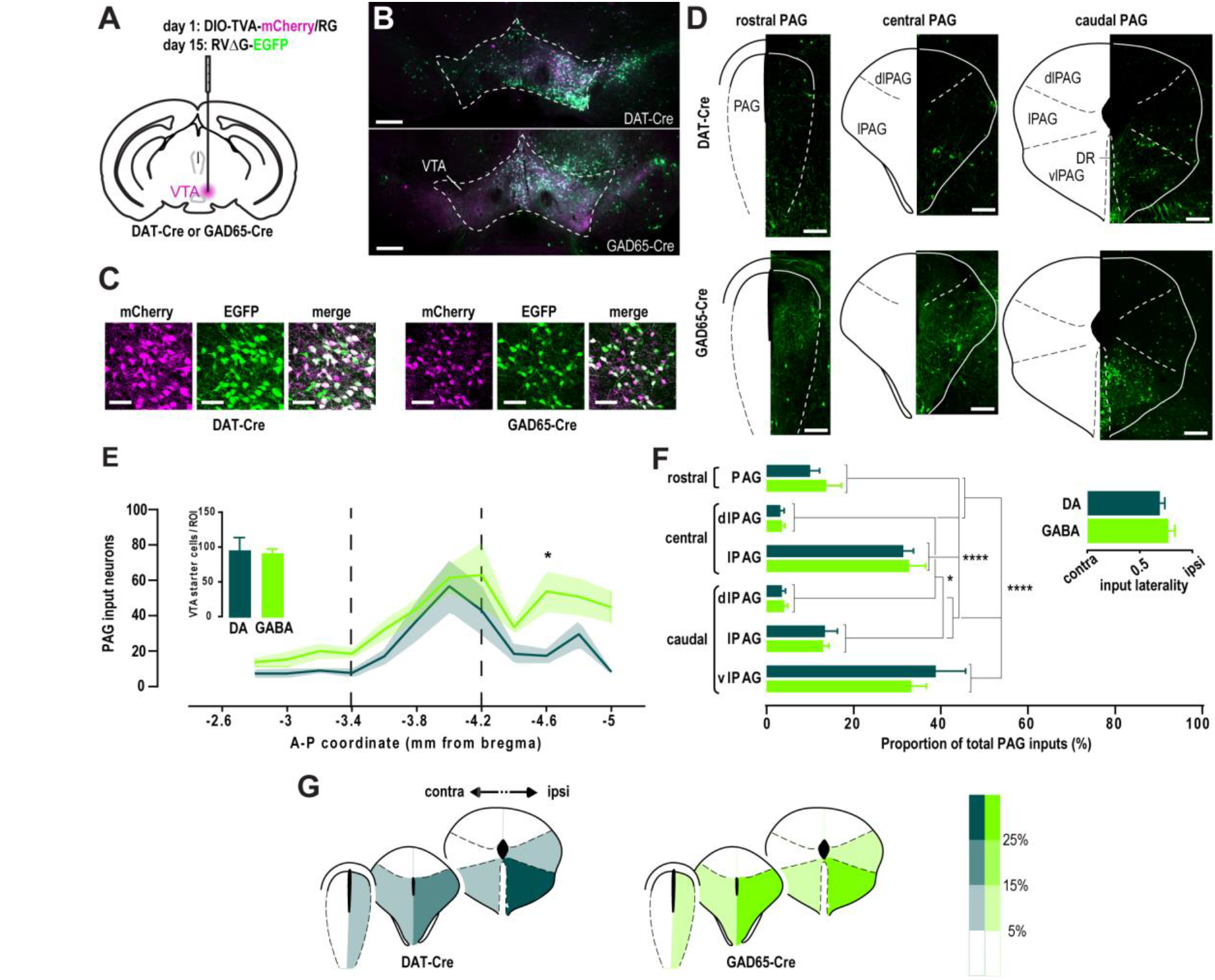
Spatial distribution of VTA-projecting PAG inputs. **A**, Schematic of the unilateral rabies injection protocol. **B-C**, Confocal images at low (B) and high (C) magnification showing the expression of TVA-mCherry (magenta) and RVΔG-EGFP (green) in the VTA seven days after the last injection. Scale bars, 500 μm (B) and 50 μm (C). **D**, RVΔG-EGFP-expressing retrogradely labeled input neurons across the rostral, central and caudal PAG segments. Scale bars, 200 Δm. **E**, Average number of inputs to VTA DA (DAT, *n* = 5) and GABA (GAD, *n* = 5) neurons along the rostro-caudal axis of the PAG (two-way ANOVA: no interaction between the cell type factor and AP coordinate factor, *F*_11,120_ = 0.7027, *p* > 0.05; main effect of cell type, *F*_1,120_ = 18.2, *p* < 0.0001; main effect of AP coordinate, *F*_11,120_ = 6.31, *p* < 0.0001; Bonferroni post-hoc test, * *p* < 0.05). Dashed lines denote the boundaries between rostral, central and caudal PAG. Inset shows the average number starter cells per ROI in DAT-Cre and GAD65-Cre mice (two-tailed *t* test: no difference between genotypes, *p* > 0.05). **F**, Relative contribution of different PAG subregions to the total inputs to VTA DA and GABA neurons (two-way ANOVA: no interaction between the cell type factor and subregion factor, *F*_5,48_ = 0.5013, *p* > 0.05; no main effect of cell type, *F*_1,48_ < 0.0001, *p* > 0.05; main effect of subregion, *F*_5,48_ = 42.85, *p* < 0.0001; Bonferroni post-hoc test, * *p* < 0.05, **** *p* < 0.0001). Inset shows the degree of lateralization of the PAG inputs (two-tailed *t* test: no difference between genotypes, *p* > 0.05). **G**, Color-coded representation of the relative input contribution of ipsilateral and contralateral PAG subregions.

### VTA DA and GABA neurons receive a similar excitatory input from the PAG

Because the PAG is a heterogeneous structure containing glutamate- as well as GABA-releasing projection neurons [19,20], we sought to investigate the nature of the VTA inputs from the PAG and test whether they exhibited a preferential targeting towards DA or GABA neurons. We thus performed whole-cell patch clamp recordings on acute VTA slices from animals transfected in the l/vlPAG with the light-sensitive cation channel ChR2 (Fig. 2A). VTA DA or GABA neurons were visually identified by the Cre-dependent transfection of the red fluorescent reporter mCherry in DAT-Cre or GAD65-Cre mice, respectively. Their neurochemical identity was confirmed by the immunohistochemical co-localization (DAT-Cre) or exclusion (GAD65-Cre) of the catecholaminergic marker tyrosine hydroxylase (TH) (Fig. 2B). Consistently with the tracing results, both DA and GABA neurons received PAG synaptic inputs with similar current amplitudes (DA vs GABA: 50.0 ± 11.2 pA vs 55.9 ± 14.2 pA) and connection rates (DA vs GABA: 40.2 ± 10.1 *%* vs 29.6 ± 5.0 %) (Fig. 2C). We chose to employ a human synapsin (hsyn) promoter-driven ChR2 in order to transfect non-selectively excitatory and inhibitory PAG projection neurons and record both excitatory inputs in the same postsynaptic neurons. Light-evoked currents were then pharmacologically blocked with glutamatergic or GABAergic antagonists in order to determine the proportion of excitatory and inhibitory afferents to VTA DA and GABA neurons (Fig. 2D). The majority of PAG inputs to either VTA DA and GABA neurons were abolished by the application of the AMPA/NMDA antagonist kynurenic acid (kyn) (DA, *n* = 16/17; GABA, *n* = 14/18), whereas the remaining kyn-resistant currents were blocked by bath application of the GABA_A_ blocker PTX (Fig. 2D-E). This suggests that PAG projections to the VTA are predominantly excitatory and do not show a preferential targeting towards either cell type. Finally, we tested whether basic electrophysiological properties of connected cells could predict the strength of the synaptic connection. We therefore analyzed the amplitude of the hyperpolarization-activated cation current *I*_h_ but found no correlation with the postsynaptic current amplitude in either VTA DA or GABA neurons. However, when we compared connected and non-connected neurons, we observed a statistically significant higher *I*_h_ current in both cell types (DA, connected vs non-connected: 91.3 ± 24.8 pA vs 45.3 ± 13.4 pA; GABA, connected vs non-connected: 40.7 ± 12.8 pA vs 19.8 ± 4.6 pA) (Fig. 2F), suggesting a higher expression of the *I*_h_-mediating HCN channels within the PAG-targeted VTA neurons. Altogether, these results describe a predominantly glutamatergic PAG-to-VTA pathway equally targeting DA and GABA neurons, and preferentially contacting neurons exhibiting a larger *I*_h_ amplitude.

**Figure 2.**
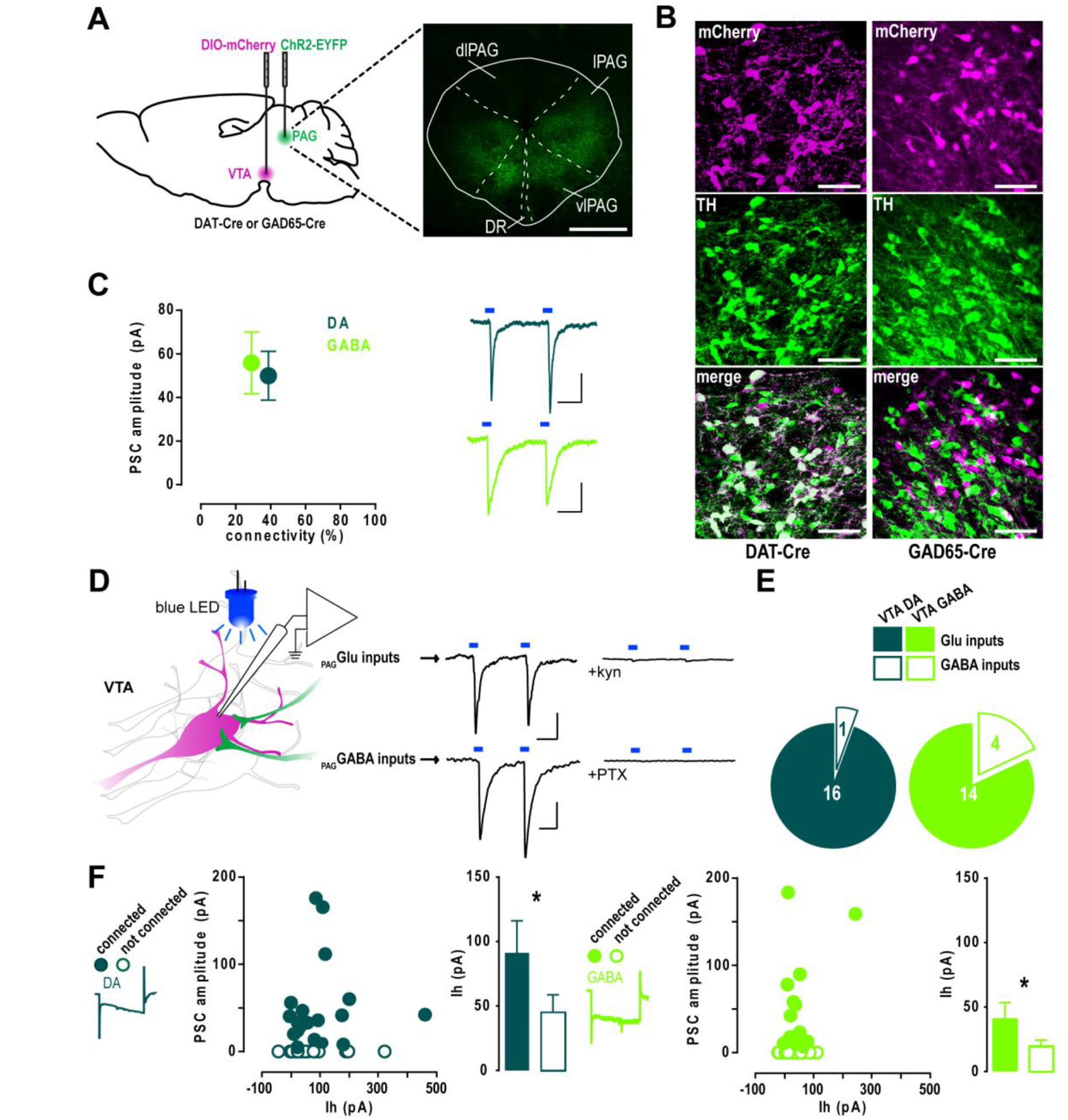
PAG afferents equally target VTA DA and GABA neurons. **A**, Left, schematic of the injection protocol for patch clamp experiments. Right, example of image of ChR2-EYFP infection in the PAG. Scale bar, 500 μm. **B**, High magnification confocal images showing the colocalization or exclusion of mCherry and TH in DAT-Cre or GAD65-Cre mice, respectively. Scale bars, 50 μm. **C**, Mean amplitude of the light-evoked postsynaptic currents in VTA DA (*n* = 47) and GABA (*n* = 62) neurons plotted against the percentage of connected neurons (Mann Whitney *U* test: no difference in amplitudes, *p* > 0.05; Fisher's exact test: no difference in connectivity, p > 0.05). Scale bars, 20 ms, 20 pA. **D**, Left, schematic of the patch clamp experiments: whole-cell recordings were performed from mCherry-expressing VTA neurons while PAG afferents inputs were light-stimulated (left). Right, excitatory currents were blocked with kynurenic acid (kyn), while kyn-resistant inhibitory currents were blocked with picrotoxin (PTX). Scale bars, 20 ms, 20 pA. **E**, Proportion of kyn-sensitive glutamate inputs and PTX-sensitive GABA inputs in VTA DA (*n* = 17) and GABA (*n* =18) neurons (Fisher's exact test: no difference between cell types, *p* > 0.05). **F**, Mean light-evoked current amplitude plotted against *I*_h_ amplitude (Spearman's rank correlation: no correlation between the variables, *r* = 0.2702, *p* > 0.05) and comparison of *I*_h_ between connected and non-connected VTA DA (left) or GABA neurons (right) (Mann Whitney *U* test: * *p* < 0.05).

## Discussion

Our anatomical tracing data indicate that PAG afferents to the VTA project towards both DA and GABA neurons, consistently with earlier observations in the rat VTA [14]. We describe here the spatial organization of the PAG inputs and provide electrophysiological evidence of functional PAG synapses similarly impinging on DA and GABA neurons of the mouse VTA.

Rabies-assisted circuit tracing has been used to obtain a brain-wide map of the afferents to neurotransmitter-defined VTA neurons [13]. This study identifies the PAG as input. Our findings expand on their results by providing a spatial profile of the location of the VTA projection neurons along the rostro-caudal and dorso-ventral axes of the PAG. We found that both VTA DA-and GABA-projecting PAG neurons were more concentrated in the central and caudal PAG, this latter segment containing a higher number of inputs to VTA GABA neurons. This comparison requires various controls. First, the absolute number of inputs depends on the size of the population of VTA DA or GABA starter neurons. Therefore, we transfected a similar number of DA and GABA starter cells in DAT-Cre and GAD-Cre mice, respectively, despite DA neurons being more frequent than GABA neurons, in a 2:1 ratio [21]. Second, distinct neuronal types might have different degrees of input convergence and different connectivity rates. VTA GABA neurons receive more convergent inputs than DA neurons, thus adding a confounding variable in the case of a starter cell-normalized comparison of the input neurons between cell types [13]. For this reason we also analyzed the number of inputs in each subregion normalized to the total PAG inputs to conclude that lPAG and vlPAG neurons provide the majority of periaqueductal efferents to VTA DA and GABA neurons.

The anatomical organization may have functional consequences. Neurons in the PAG have been proposed to subserve different behavioral functions – from opioid analgesia to modulation of autonomic responses and defensive behaviors, depending on their columnar localization. For instance, two distinct types of opioid-induced and opioid-independent pain suppression mechanisms have been proposed to arise from the stimulation of the ventrolateral or lateral columns of the PAG, respectively [2]. Analogously, two antagonistic threat responses, the freezing and flight behaviors, were recently shown to be promoted by neurons within vlPAG and dl/lPAG columns, respectively [22]. In light of these observations, it is tempting to speculate that our results on the PAG inputs distribution pattern might reflect their involvement in opioid analgesia and freezing responses. However, further experiments are needed to test this hypothesis and to determine the physiological relevance of this projection to the VTA.

Both glutamate- and GABA-releasing output neurons have been described in the PAG [20,23]. We therefore tested whether there would be a neurotransmitter-specific organization these PAG projections to either DA or GABA neurons of the VTA. However, our results did not support this hypothesis, suggesting instead that the PAG provides an equal synaptic input to both cell-types. Since the electrophysiological markers classically used to distinguish between VTA DA and GABA neurons (e.g. action potential width, capacitance measures and presence of a hyperpolarization-activated *I*_h_ current) are not reliable, in light of their variability and overlap between the two cell types [24,25], we chose to employ a Cre-dependent genetic approach in order to unequivocally identify DA and GABA neurons in DAT-Cre and GAD65-Cre mice, respectively. In accordance with previous observations, both identified DA and GABA neurons exhibited a variable range of *I*h amplitudes [24]. Additionally, our results suggest that PAG projections preferentially target neurons exhibiting larger *I*h currents. It is plausible that these differences reflect a higher expression, or a change in the subunit composition of the hyperpolarization-activated cyclic nucleotide-gated (HCN) channels, which mediate the *I*h currents and have been implicated in the reward system adaptations induced by drugs of abuse [26,27].

Our findings indicate that PAG GABA neurons constitute a small fraction of the projection to the VTA targeting the VTA neurons via GABA_A_ transmission. These results rely on non-selective expression of ChR2 under the hsyn promoter in all PAG projection neurons. However, the possibility that this promoter might differentially drive the expression of ChR2 among cell-types seems unlikely. In comparison, a previous ultrastructural analysis of PAG axons within the VTA reported that only 8% immunostained for GABA [14], although the authors note that immunolabeling of electron microscopy samples may be prone to false negatives and, consequently, the actual contribution of GABA neurons to the PAG input is likely to be higher. Despite this possible underestimation, our electrophysiological results provide further support to the observation of a small GABAergic participation in the PAG-to-VTA pathway. It should be noted, however, that our findings do not take into account the potential presence of inhibitory synapses expressing only the metabotropic GABAB receptor, as it was recently described for the NAc projections to VTA DA neurons [28].

## Conclusion

Neurons of the VTA are known to be a major target of addictive drugs [29], and their activity has been associated with both rewarding and aversive behaviors [12,30]. Similarly, PAG neurons have been implicated in the expression of opioid-induced analgesia and in the modulation of fear responses [1]. The anatomical and electrophysiological data presented here thus expand on the existing knowledge of the connectivity between VTA and PAG neurons, providing a substrate for future investigations on their interaction during motivationally relevant behaviors.

## Acknowledgements

We thank all members of the Lüscher laboratory for stimulating discussions and valuable comments on the manuscript.

